# RL mechanisms of short-term plasticity of auditory cortex

**DOI:** 10.1101/093138

**Authors:** Elena Krugliakova, Alexey Gorin, Anna Shestakova, Tommaso Fedele, Vasily Klucharev

## Abstract

The decision-making process is exposed to modulatory factors, and, according to the expected value (EV) concept the two most influential factors are magnitude of prospective behavioural outcome and probability of receiving this outcome. The discrepancy between received and predicted outcomes is reflected by the reward prediction error (RPE), which is believed to play a crucial role in learning in dynamic environment. Feedback related negativity (FRN), a frontocentral negative component registered in EEG during feedback presentation, has been suggested as a neural signature of RPE. In modern neurobiological models of decision-making the primary sensory input is assumed to be constant over the time and independent of the evaluation of the option associated to it. In this study we investigated whether the electrophysiological changes in auditory cues perception is modulated by the strengths of reinforcement signal, represented in the EEG as FRN.

We quantified the changes in sensory processing through a classical passive oddball paradigm before and after performance a neuroeconomic monetary incentive delay (MID) task. Outcome magnitude and probability were encoded in the physical characteristics of auditory incentive cues. We evaluated the association between individual biomarkers of reinforcement signal (FRN) and the degree of perceptual learning, reflected by changes in auditory ERP components (mismatch negativity and P3a). We observed a significant correlation of MMN and valence - dFRN, reflecting differential processing of gains and omission of gains. Changes in P3a were correlated to probability - dFRN, including information on salience of the outcome, in addition to its valence.

MID task performance evokes plastic changes associated with more fine-grained discrimination of auditory anticipatory cues and enhanced involuntary attention switch towards these cues. Observed signatures of neuro-plasticity of the auditory cortex may play an important role in learning and decision-making processes through facilitation of perceptual discrimination of valuable external stimuli. Thus, the sensory processing of options and the evaluation of options are not independent as implicitly assumed by the modern neuroeconomics models of decision-making.

## 1. Introduction

Decision theory assumes that individuals’ choices are driven by values, attached to prospective outcomes. In order to evaluate expected values (EV) of options, individuals estimate the magnitude and probability of outcomes (Bandura, 1977; Von Neumann and Morgenstern, 1944). Neural correlates of EV (including magnitude and probability of outcomes) have been widely investigated during last two decades (see Glimcher et al., 2009 for a review). In neurobiological models of decision-making (Rangel et al., 2008), the primary sensory input is assumed to be independent of the evaluation of the option associated to it. In particular, while the decision-making process is exposed to modulatory factors, the sensory input, which is a basis for this process is considered to be constant over time. However, experience-induced plasticity is vital for human brain and provides possibility to learn and adapt to constantly changing environment. Moreover, neuroplastic changes are observed not only during ontogenesis but long after the developmental period (Buonomano and Merzenich, 1998). For example, in the motor cortex and sensory cortices of different modalities, learning leads to the increase in the number of neurons that represent the learned stimuli or induces the spatial rearrangement in neurons populations topography (Nudo et al., 1996; Recanzone et al., 1992a, 1992b). An experience-driven improvement in quality of stimuli perception is called *perceptual learning* and can be used to explore plasticity in sensory cortices (Gilbert et al., 2001). Numerous ERP studies have shown training-induced neuroplastic changes in auditory information processing (Atienza et al., 2005; Kujala and Näätänen, 2010; Shtyrov et al., 2010) which could be explained by reorganization of neuronal populations and changes in sensitivity and processing of relevant information. In particular, frequency discrimination training is shown to enlarge the cortical representation of the corresponding frequencies (Recanzone et al., 1993) as more neurons become responsive to trained frequencies.

In auditory sensory processing, brain plasticity is represented by the mismatch negativity (MMN). MMN is evoked by the presentation of an oddball or deviant stimuli (150–250 ms after deviant onset), embedded in a stream of repeated stimuli, the standards (Näätänen et al. 2007). The MMN component can be elicited out of the focus of attention and is thought to be the result of a pre-attentive process able to detect alterations in a regular sound sequence (Näätänen, 1990; Winkler et al., 1996).Changes in the amplitude and latency of MMN seem to reflect the more fine-grained stimulus discrimination indicating sensory cortex neuroplasticity. Initial poor identification of differences between deviant and standard stimuli as well as inaccuracy of performance is correlated with a low-amplitude MMN, while following an active learning to discriminate deviant pattern results in emerging MMN activity (Cheour et al., 2002; Näätänen et al., 1993a; Novak et al., 1990; Sams et al., 1985; Tiitinen et al., 1994). A number of magnetoencephalographic (MEG) studies showed that training-dependent MMN is generated in the auditory cortex (Alho et al., 1996; Tervaniemi et al., 2001). Furthermore is was demonstrated that presence of changes in MMN amplitude not only right after discrimination training, but also several days later (Atienza et al., 2002, 2005; Kraus et al., 1995; Menning et al., 2000; Tremblay et al., 1998), suggesting a training-dependent long-term effect on pre-attentive sensory processing in the auditory cortex. Overall, the MMN amplitude was found to be enhanced and/or its latency shortened as a consequence of training or due to long-term experience (Cheour et al., 1998; Kraus et al., 1995, 1996; Tervaniemi et al., 2001; Winkler et al., 1999). Thus, previous studies have robustly demonstrated that the training-induced change of the MMN amplitude seem to be a reliable marker of experience-induced neuroplasticity.

Training-induced enhancement in MMN might be accompanied by increase in P300 – a positive ERP component, known to be induced by infrequent stimulus presentation (Sutton et al., 1965). P300 consists of two sub-components: P3a with latency 230–300 ms and fronto-central distribution, and classic P300 (P3b), with latency 300–400 ms and parietal distribution. In particular P3a, which reflects attentional reorientation to salient, task irrelevant cues (Escera et al., 1998; Wetzel et al., 2011) has been linked to both short- and long-term plasticity changes as a result of auditory training (Atienza et al.2004; Draganova et al. 2009; Uther et al. 2006). The P3a is assumed to reflect the orienting response associated with attentional reorientation to stimuli, salient but irrelevant for task (Escera et al., 1998; Wetzel et al., 2011), it is associated with executive functions (Fjell et al., 2009; Light et al., 2007) and possibly working memory encoding (Bledowski et al., 2004). P3a response has been linked to both short- and long-term plasticity changes as a result of auditory training (Atienza et al.2004; Draganova et al. 2009; Uther et al. 2006). Changes in P3a properties are associated with an involuntary attention shift to the deviant tone due to familiarization mediated by top-down process resulting in a rapid learning (Draganova et al., 2009).

Thus, the MMN and the P3a components has been successfully used as an electrophysiological measure of plasticity in the auditory system, which to our knowledge has not been taken into account in neuroeconomics models of EV processing. More specifically, we hypothesized that building association between auditory stimuli physical properties (such as frequency and intensity) and components of EV (magnitude and probability) might induce enlargement in auditory ERP components - MMN and P3a, measured by means of classical passive oddball paradigm. Changes in MMN would indicate perceptual learning, associated with experience-induced brain plasticity of the auditory cortex and resulting in more fine-grained discrimination of auditory stimuli (bottom-up process) (Draganova et al., 2009). Whereas changes in P3a would indicate reallocation of attention to relevant stimuli, guided by prefrontal cortex (top-down process) (Polley et al, 2006). The prediction of training-induced changes in sensory perception, based on effectiveness of learning, could help to understand how fundamental mechanisms of reinforcement learning and sensory plasticity may be interconnected in behavioral adaption.

One of the most studied electrophysiological correlates of reinforcement learning is the ERP component known as feedback related negativity (FRN), a frontocentral negative ERP component, occurring 240–340 ms after feedback onset. The FRN is believed to represent an alerting signal, following unexpected and/or unfavourable outcomes that underlies learning and performance monitoring (Holroyd and Coles, 2002; Van Meel et al., 2005; Montague and Berns, 2002; Montague et al., 2004; Sambrook and Goslin, 2015b). An influential theory (Holroyd & Coles, 2002) suggests that FRN, codes reward prediction error (RPE). RPE reflects a discrepancy between the obtained outcome and expected outcome. Unexpected and/or unfavourable outcomes (i.e. monetary loss) produce negative RPEs, whereas unexpected favorable outcomes (i.e. monetary gain) result in positive RPEs. The FRN is known to be strongly affected by valence of feedbacks – it is enhanced by unfavourable outcomes (Miltner et al., 1997; Sambrook and Goslin, 2015a). EEG and fMRI researches stated a causal role of dopaminergic system in FRN generation in the cingulate cortex (for review, see Walsh and Anderson, 2012). An EEG study showed that FRN is a better index of negative RPE as compared to positive RPE (Gehring and Willoughby, 2002). This sensitivity to valence of the outcome constitutes the core argument proving that FRN might be an encoder of RPE-sign. Importantly negative RPE is generated when outcomes (a monetary loss or omission of monetary gain) are worse than predicted (i.e, Holroyd and Coles, 2002; Luu et al., 2000). Overall, neuroimaging studies suggest that FRN is more sensitive to probability of outcomes than to the magnitude of outcomes (Walsh and Anderson, 2012), despite some evidences that outcome magnitude exerts a modulatory effect on FRN (Sambrook and Goslin, 2015a).

The monetary incentive delay task is an elegant tool to study different stages of reinforcement learning, from reward anticipation to its delivery (Knutson et al., 2000, 2005). It can be used to delineate neural mechanisms of performance monitoring during behavioral acts with different EVs and RPEs. By introducing incentive cues, signaling both magnitude and probability of prospective outcomes it is possible to study their effects on neural activity, associated with feedback evaluation (Knutson et al., 2005). Initially, MID task was used in fMRI studies of reward processing (Knutson et al., 2000). Later EEG and MEG studies have started to employ MID to study neural dynamics of reward processing (Broyd et al., 2012; Doñamayor et al., 2012; Thomas et al., 2013). To our knowledge, none of the previous studies investigates the association between the FRN characteristics, registered in MID task delivery phase, and consequent changes in perception of incentive cues, marking the onset of the MID anticipatory phase. In classic MID task, visual stimuli such as circles, squares and triangles were utilized as an incentive cues coding probabilities and magnitudes of outcomes. We developed an auditory version of MID task, which was used to study auditory perceptual learning. Sounds of different frequencies were used as incentive cues signaling prospective gain’s probabilities and magnitudes.

It has been shown, that the FRN amplitude can serve as an individual index of learning effectiveness. There is strong evidence approving hypothesis that the stronger the FRN is, the better the participant learns (Luft, 2014; Luft et al., 2013). In our study, correlation of FRN values, modulated either by probability or magnitude of outcome, with changes in auditory stimuli perception after MID task performance would provide evidence of interplay between reinforcement learning mechanisms and induced plasticity of sensory processing.

Overall, in our work we tested two hypothesis: (a) repetitive MID task will induce changes in MMN and P3a and these changes will depend on the expected value assigned to tone used as acoustic cue (b) these changes can be predicted by the size of FRN, elicited during feedback presentation in MID task.

## 2. Material and Methods

### 2.1 Subjects

Seven subjects (4 women, 22±2 y.o.) participated in the behavioural pilot experiment. Forty-two subjects (20 women, 23±4 y.o.) participated in EEG experiment, in which both behavioural and electrophysiological data were collected. Data from five additional subjects were excluded due to excessive EEG artefacts. All subjects were right-handed, with normal or corrected-to-normal vision and without history of psychiatric or neurological disorders. The study was approved by the local Ethics Committee and all participants gave written informed consent prior to their participation.

Auditory stimuli

Three sinusoidal tones (523, 1046 and 1569 Hz with fundamental frequency corresponding to C5 of the Western musical scale, intensity 70 dB) were used in the oddball paradigm as the standard stimuli (Fig. 1).

**Figure 1.**
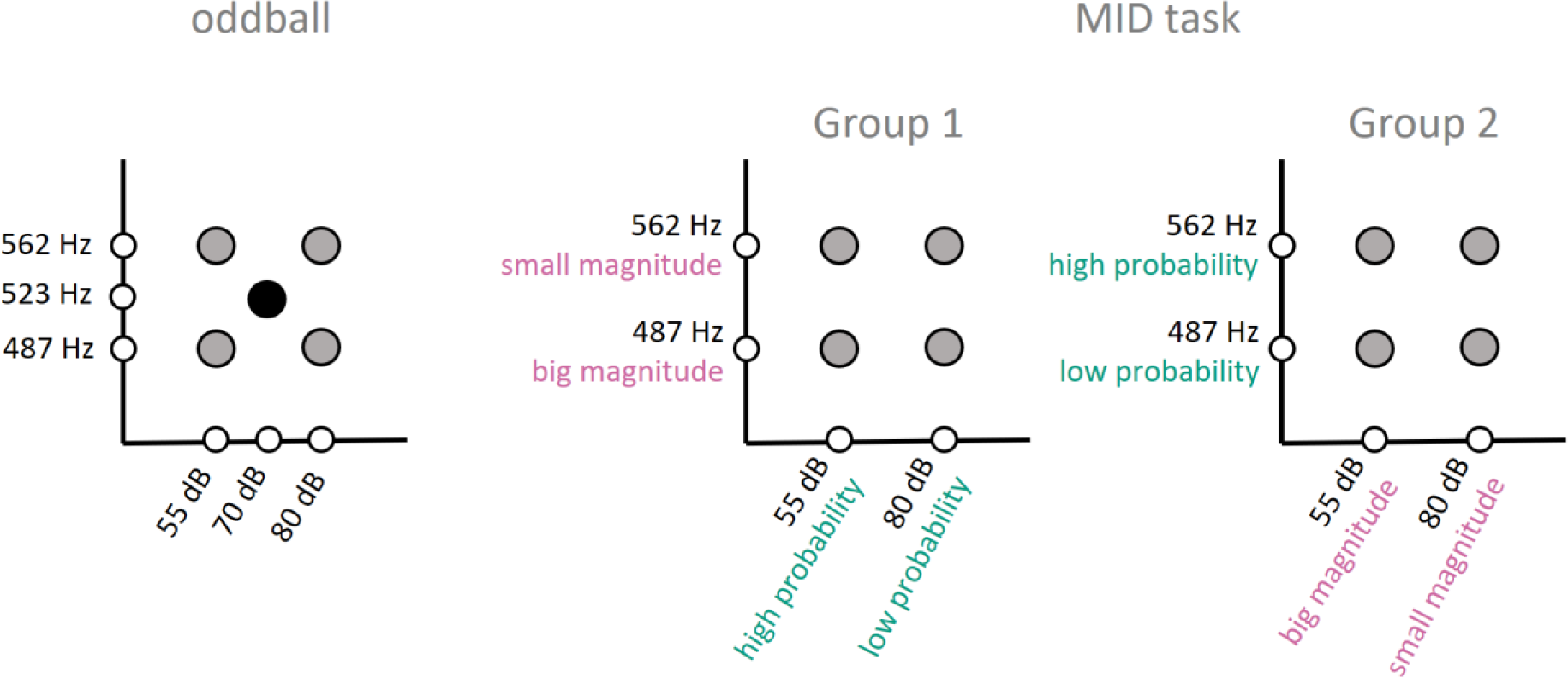
Physical parameters of acoustic stimuli. The left scheme depics the stimuli used in the oddball paradigm with deviants in gray dots and standard in black dot. Middle and right schemes show the encoding of gain magnitude and probability into frequency and intensity of the acoustic stimuli in two esperimental groups.

Deviant tones (Fig. 1, left scheme) differed from standard both in frequency and intensity according to a probabilistic design (an increment or decrement were equally probable) which resulted in four distinct deviants. Deviants differed from standards in frequency by + 10/8 and - 10/8 semitones of the Western musical scale (fundamental frequencies 562 Hz for the higher and 487 Hz for the lower deviant tones). The intensity of the deviants was either smaller or bigger than the standard (70 dB) by 15 dB and 10 dB, respectively (55 and 80 dB) (Fig. 1, left scheme). All tones had a duration of 200 ms (including 5 ms rising and falling times). Stimuli were generated with PRAAT software.

The same four deviant oddball stimuli were also used as acoustic cues in the auditory MID task. Acoustic cues signaled high or low prospective reward probability (0,80 and 0,20, correspondingly) and high or low prospective reward magnitude (4 or 20 rubles, correspondingly), as illustrated by Fig. 1 (middle and right schemes). We used two levels of frequency and two levels of intensity to encode prospective reward probability and magnitude. The probability and magnitude of reward were encoded differently in two experimental groups. In the Group 1 (n = 24) outcome magnitude was encoded by the intensity of the acoustic cue, while the gain probability was encoded by its’ frequency. In the Group 2 (n = 25) the encoding of gain magnitude and gain probability was reversed.

### 2.2 Study design

The primary goal of this study was to investigate the effect of MID task on perception and processing of auditory stimuli used as incentive cues. For this purpose, we designed an experiment consisting of two types of task presented in two successive days.

*Day 1.* At the beginning of each experiment, the ability of participants to discriminate auditory stimuli was tested during a recognition test. After the recognition test, participants performed the first session of passive oddball. Next, participants performed the first session of MID task. Prior to MID task, probe structure and meaning of each acoustic cue were explained to participants.

*Day 2.* Approximately at the same day time, participants performed the second MID task session and the second oddball. Both MID task and oddball were analogous across two experimental days.

#### Recognition test

Recognition test was designed to ensure that participants were able to discriminate acoustic cues coding expected values. The participants were instructed to press a button corresponding to the delivered sound. The sound descriptions and target buttons were displayed on the screen (i.e. high loud sound, button 1, etc.) during the task. Participants received positive and negative visual feedbacks to facilitate learning. The EEG session started when the subject successfully identified 8 out of 10 consecutive sounds. On average, participants made more mistakes in frequency identification (5.14±1.26; S±SEM) as compared to intensity identification (2.00±0.51) and as compared to mistakes in simultaneous identification of frequency and intensity of sounds (1.90±0.64).

#### Auditory MID task

During the auditory MID task (Fig. 2), participants were exposed to acoustic cues encoding prospective gain magnitude (4 or 20 rubles) and probability of win (0,80 and 0,20). After a variable anticipatory delay period (2000-2500 ms), participants responded by single button press immediately after the presentation of a visual target (white square) (Fig. 2). After a short delay, subsequent feedback notified subjects whether they had won money together with their cumulative total outcome (2000 ms).

**Figure 2.**
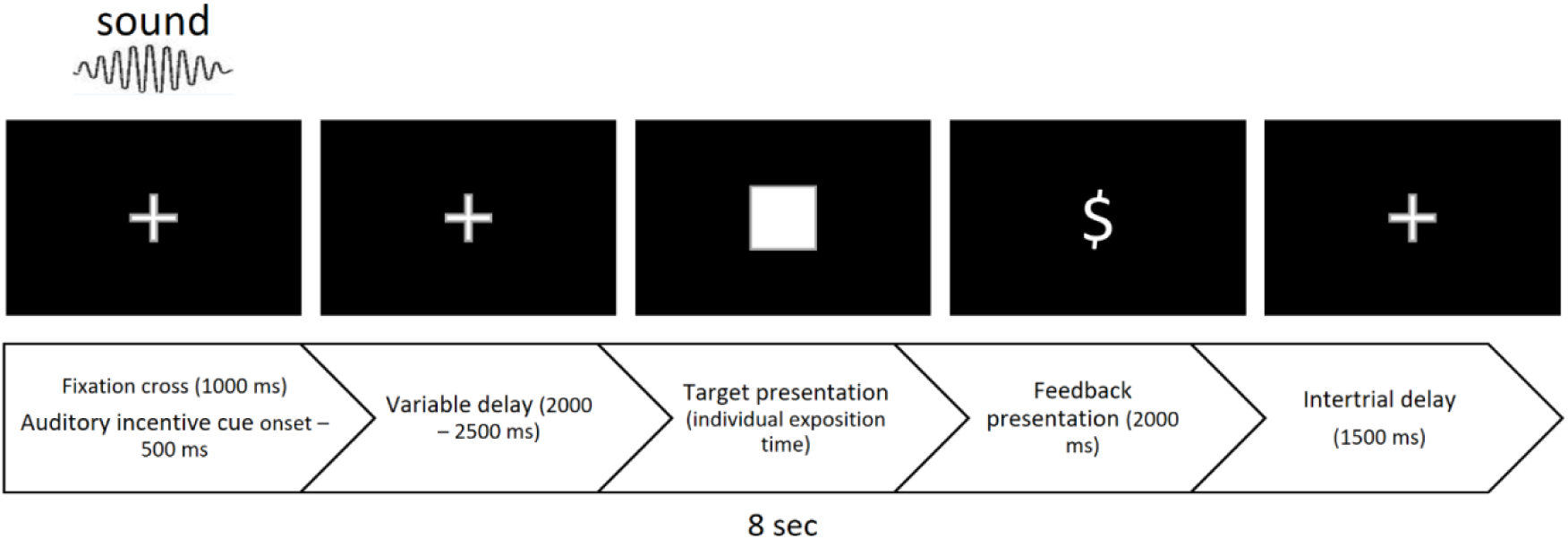
Trial scheme of auditory MID task.

Overall, outcomes were positive (gain 4 or 20 rubles) or negative (omission of gain 4 or 20 rubles). Probability of win was manipulated by altering the average target duration through an adaptive timing algorithm that followed subjects’ performance, such that they would succeed on 80% of high-probability trials and 20% of low-probability trials overall.

At the beginning of the task, duration of the target was based on reaction times collected during the training session. Importantly, prior to the MID task participants were instructed what acoustic cues corresponded to which probabilities and magnitudes of outcomes.

On average, duration of target was set to 276±29 ms for trials with high gain probability and 189±26 ms for those with low gain probability. The reward feedback was presented an average of 58±4 trials out of 76 in case of 80% gain probability and an average of 13±3 trials out of 76 in 20% gain probability trials. On average by the end of the game subjects earned 854±76 rubles.

#### Oddball paradigm.

To record MMN before and after MID training, subjects participated in the passive oddball paradigm. Infrequent deviant tones were randomly interspersed with a standard tone presented with a probability (P_std_) of 0.8 (Fig.1, b) and with a 800±100 ms onset-asynchrony (SOA). Each deviant type (Dev_I1F1_, Dev_I1F2_, Dev_I2F1_, Dev_I2F2_) was presented as every fourth, fifth or sixth tone with the same probability (P_dev_ = 0.2/4 = 0.05). Two successive deviants were always of different type (e.g., Dev_I2F2_ – Std – Std – Std – Dev_I1F1_ – Std – Std – Std – Std – Std – Dev_I2F1_ – Std – Std – Std – Std – DevI1f1 – Std –...). The stimuli were delivered in 30-min sequences: 2400 tones per sequence, each of the 4 deviants was presented 120 times. All sequences started with four successive standards.

### 2.3. Analysis of behavioral results

Reaction time (RT) on each trial type was averaged for each individual, grand averaged and subjected to mixed four - way repeated measures analyses of variance (ANOVA) with *Group* as a between-subject variable (group 1 vs. group 2) and *Session* (MID - session 1 vs. MID - session 2), *Magnitude* (low magnitude vs. high magnitude) and *Probability* (20% vs. 80%) as within-subject variables.

### 2.4. EEG data acquisition

EEG data were recorded using 28 active electrodes (Brain Products GmbH) at a sampling rate of 500 Hz, according to the extended version of 10–20 system: Fp1, Fp2, F3, F4, C3, C4, P3, P4, O1, O2, F7, F8, T7, T8, P7, P8, Fz, Cz, Pz, Oz, FC1, FC2, CP1, CP2, FC5, FC6, CP5, CP6. Active channels were referenced against the mean of two mastoids electrodes, in order to display the maximal FRN response at frontal electrode sites. The electrooculogram (EOG) was recorded with electrodes placed at the outer canthi and below the right eye. Data was acquired with a BrainVision actiCHamp amplifier (Brain Products GmbH) and sampled at 500 Hz. Impedance was confirmed to be less than 5 kΩ in all electrodes prior to recordings.

### 2.5. EEG data analysis

EEG signals were pre-processed with BrainVisionAnalyzer 2.1 (Brain Products GmbH). The EEG was filtered offline (passband 1–30 Hz, notch filter – 50 Hz), then ICA-based ocular artifacts correction was performed. After manual raw-data inspection for remaining artefacts, data were segmented to epochs of 600 ms including a 100 ms pre-stimulus. Each trial was baseline corrected to an average activity between –100 and 0 ms before stimulus onset. Epochs including voltage changes exceeding 75 mV at any channel were omitted from the averaging. Epochs were separately averaged for different trial types. Averaged ERP waveforms were computed within each subject and condition with a minimum number of 15 trials per condition.

A time windows chosen for statistical analysis of ERP components were based on visual inspection of the grand-average waveforms and previous research on the relevant components. ERP components were defined either like local maxima (P3a) or local minimum (MMN and FRN). Once a peak was identified, the amplitude over a window ±20 ms around this peak was averaged individually and then grand averaged across participants.

#### 2.5.1. Auditory MID task EEG data analysis

In order to disentangle the influence of two RPE modulators, we processed feedback-locked visual ERP in two different ways: pooling ERPs for expected (likely) and unexpected (unlikely) outcomes, irrespective to magnitude, and pooling ERPs for high (20 rub) and low (4 rub) magnitude, irrespective of probability. ERPs obtained during the first and the second session were pooled together. As a result, we obtained 4×2 different types of waveforms. This procedure also helped to increase the number of trials averaged for each type of feedback, because of a large difference in number of trials for expected and unexpected outcomes. Peak amplitudes of components were quantified as the average amplitude around (±20 ms) the local minimum occurring within the timeframe of interest (270-350 ms on electrode Fz) post stimulus-onset. Timeframes of interest were the same for all eight types of feedback. For probability-pooled ERPs we averaged 27±6 unlikely positive outcomes (gain in case of 20% probability of success), 110±14 likely positive outcomes (gain in case of 80% probability of success), 28±7 unlikely negative outcomes (miss in case in case of 80% probability of success) and 110±19 likely negative outcomes (miss in case of 20% probability of success). For magnitude-pooled ERPs we averaged 67±5 low magnitude positive outcomes (4 rub gains), 84±10 high magnitude positive outcomes (20 rub gains), 71±8 low magnitude negative outcomes (4 rub misses) and 68±7 high magnitude negative outcomes (20 rub misses). ERPs obtained during the first and the second session were pooled together.

For probability-pooled ERPs we calculated ERPs (27±6 trails) to unlikely positive outcomes (gain, p=0,20), ERPs (110±14 trials) to likely positive outcomes (gain, p=0,80), ERPs (28±7 trails) to unlikely negative outcomes (miss, p=0,80) and ERPs (110±19 trials) to likely negative outcomes (miss, p=0,20). For magnitude-pooled ERPs we calculated ERPs (67±5 trials) to small positive outcomes (4 rub), ERPs (84±10 trials) to big positive outcomes (20 rub), ERPs (71±8 trials) to low negative outcomes (misses of 4 rub) and ERPs (68±7 trials) to high negative outcomes (misses of 20 rub). ERPs obtained during the first and the second sessions were pooled together. Mixed four - way repeated measures ANOVAs with *Group* as a between-subject variable (group 1 vs. group 2) and *Valence* (gain vs. miss), *Probability* (unlikely vs. likely) and *Electrode* (Fz vs. Cz vs. Pz) as within-subject variables were conducted for FRN amplitudes derived from probability-pooled ERPs. Mixed four - way repeated measures ANOVAs with *Group* as a between-subject variable (group 1 vs. group 2) and *Valence* (gain vs. miss), *Magnitude* (low magnitude vs. high magnitude), *Electrode* (Fz vs. Cz vs. Pz) as within-subject variables were conducted for FRN amplitudes derived from magnitude-pooled ERPs.

In addition to analysis of FRN amplitudes, we calculated differential FRN (dFRN). These values was further used as individual biomarkers of reinforcement learning effectiveness. *Valence dFRN* (FRN to all positive outcomes (gain) *minus* FRN to all negative outcomes (omission of gain). *Probability dFRN* and *Magnitude dFRN* were calculated similar to Sambrook and Goslin, 2015a. By subtracting waveforms for gains and misses with the same size of RPE, we obtained difference waveforms reflecting differences in processing feedback valence in case of small RPE and big RPE. Then, we subtracted obtained difference waveforms for small RPE from waveforms for big RPE. Thus, the overall scheme of *probability dFRN* calculation was as follows: ((unexpected misses – unexpected gains) – (expected misses – expected gains)), irrespective to magnitude of outcome. For *magnitude dFRN* the calculation scheme was similar ((big misses – big gains) – (small misses – small gains)). Amplitudes of *valence dFRN, probability dFRN* and *magnitude dFRN* were detected with the same procedure used for amplitudes of FRN. We conducted two-way mixed model ANOVA with dFRN amplitudes as dependent variable, with *Group* as a between-subject variable (group 1 vs. group 2) and *FRN type* (valence vs. probability vs. magnitude) as within-subject variable. Timeframes of interest for the three dFRN were the same as for FRN. This procedure allowed us to compare the sensitivity of FRN and dFRN to valence and to components of EV. For dFRN values, we conducted two-way mixed model ANOVA with dFRN amplitudes as dependent variable, with *Group* as a between-subject variable (group 1 vs. group 2) and *FRN type* (valence vs. probability vs. magnitude) as within-subject variable.

#### 2.5.2. Oddball EEG data analysis

To study experience-induced plastic changes we analyzed MMN and P3a components before and after the training in MID task. For the oddball analysis, data were segmented for five types of trials: standard stimulus and four types of deviants. MMN and P3a were derived by subtraction of the averaged response to standard stimulus from the averaged response to each type of deviant stimulus. The MMN peak amplitude was identified as the most negative peak in the obtained difference curve occurring at 80–250 ms post stimulus-onset on electrode Cz (Näätänen, Paavilainen, Rinne, & Alho, 2007). The P3a peak amplitude was identified as the most positive peak of difference curve occurring at 180-300 ms post stimulus-onset on electrode Fz (Seppänen et al., 2012).

Mixed five - way repeated measures ANOVAs with *Group* as a between-subject variable (group 1 vs. group 2) and *Session* (MID - session 1 vs. MID - session 2), *Magnitude* (4 rub vs. 20 rub), *Probability* (20% vs. 80%), and *Electrode* (Fz vs. Cz vs. Pz) as within-subject variables were used to assess dependence of MMN and P3a amplitudes on two components of EV as well as day of experiment and electrode, where data were recorded.

In all repeated measures ANOVA analyses, significant interactions were further decomposed with simple effect tests (Howell and Lacroix, 2012; Stevens, 1991). The level of significance was set to p < 0.05. P-values reported for ANOVAs analyses were adjusted with the use of the Greenhouse– Geisser correction. All statistical analyses were performed using the Matlab 2015a and SPSS software package (22.0).

### 2.6. dFRN-MMN Correlation

We analyzed whether the changes in MMN and P3a can vary as a function of dFRN amplitude, registered in MID task. For this purpose, we conducted two sets of correlation analysis, one for MMN and one for P3a. To estimate changes across oddball sessions in MMN and P3a we subtracted peak values for Session 1 from values for Session 2. Thus, the more negative (positive) value, the bigger MMN (P3a) was on the second day.

To quantify the relationship between changes in amplitudes of MMN and P3a (oddball) and valence, probability and magnitude dFRNs (MID), we calculated Spearman correlation between these two classes of variables. Cook’s distance was used to identify outliers. Cases with Cook’s distance bigger 4/n were excluded from the further analysis (Bollen and Jackman, 1985).

## 3. Results

### 3.1. Behavioral results: auditory MID

Mean reaction time (RT) of participants was 232±25 ms. RT in each trial type was averaged individually for MID - session 1 and MID - session 2 (Fig. 3). A repeated measures ANOVA revealed significant main effects of Group [F(1, 48) = 5.708, p = 0.021, η2p = 0.275]: we observed a longer average RT in the Group 1 (221±5) as compared to the Group 2 (204±5). Probability and valence of the expected outcome significantly modulated RTs: factors Probability [F(1, 48) = 135.632, p < 0.001, η2p = 0.739] and Magnitude [F(1, 48) = 18.209, p < 0.001, η2p = 0.275]. In average, participants were faster in trials with low probability of positive outcomes (202±4) as compared to those with high probability (222±3). RT was faster in trials with large magnitude of expected gains (210±3) as compared to trials with small magnitude (215±4). No significant interactions between factors were observed.

**Figure 3.**
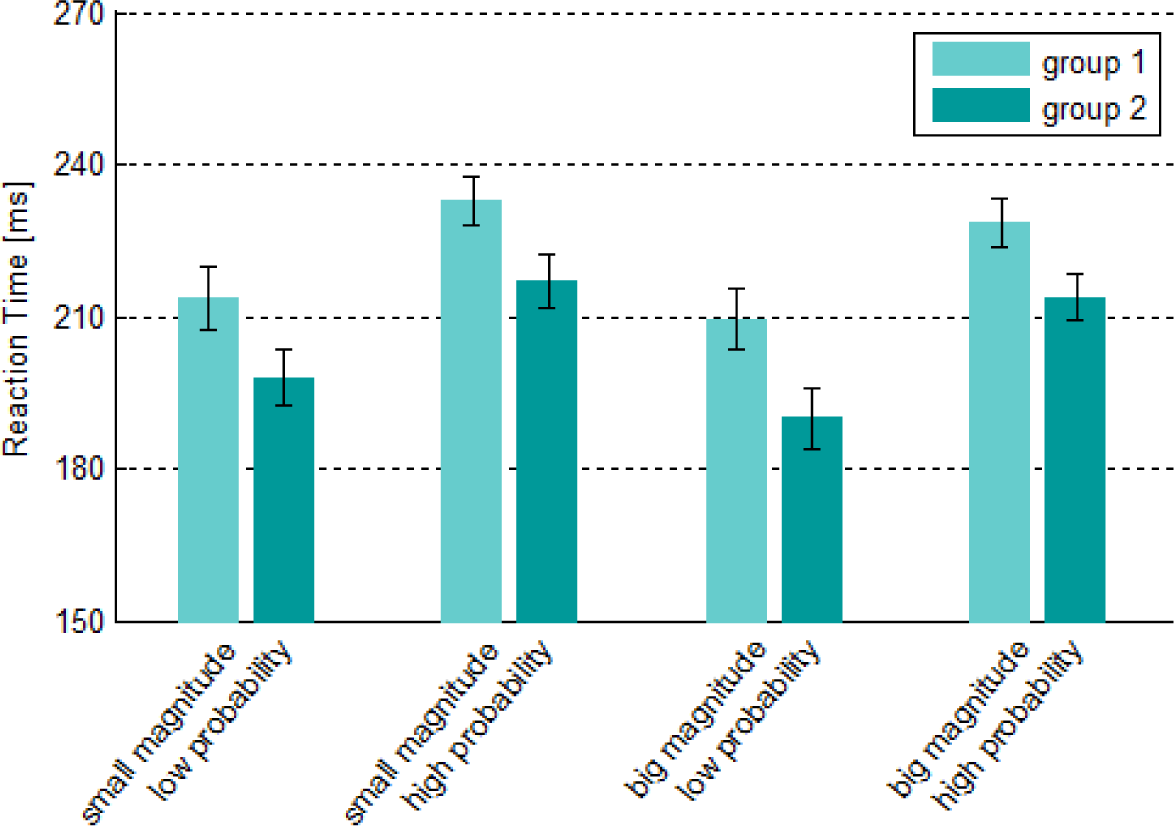
RTs for different types of trials in two experimental groups (light blue – group 1, green – group 2).

### 3.2. Electrophysiological results: auditory MID

Fig. 4 presents eight different types of feedback-locked visual ERP waveforms recorded in MID at Fz. In all conditions feedbacks are followed by FRN as a negative deflection around 300 ms. For both experimental groups FRN was stronger for negative outcomes than for positive outcomes: the smallest FRN was evoked by unexpected gains; while small negative outcomes resulted in the largest FRN (Fig. 4).

**Figure 4.**
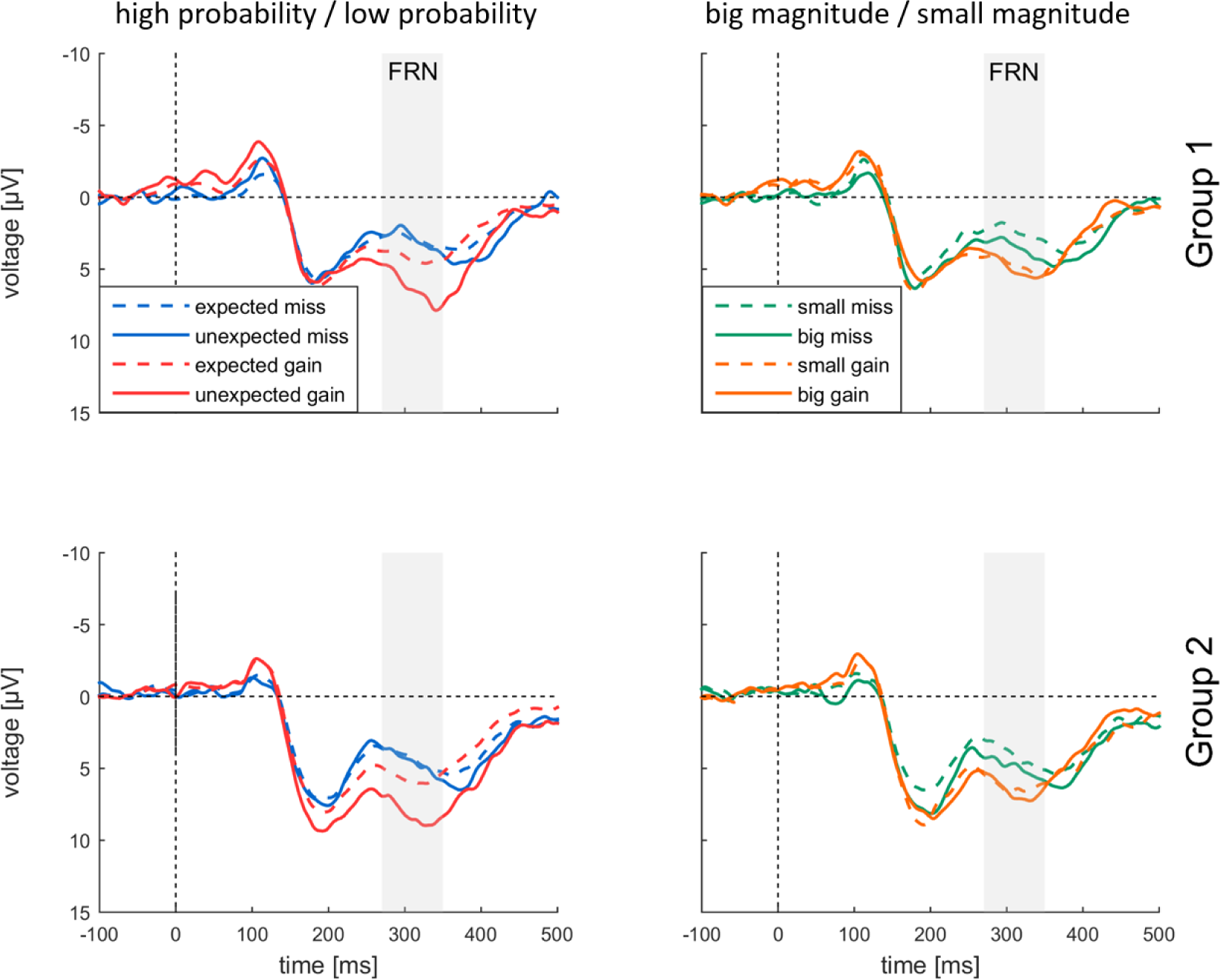
Grand-averaged visual ERP waveforms superimposed for eight types of feedbacks (averaged across two MID task sessions), as a function of probability (left part) or magnitude (right part). FRN component (270 – 350 ms) is highlighted by gray shadings).

Main effect of *Valence* [*F*_(1,40)_ = 25.892, *p* < 0.001, η^2^_*p*_ = 0.393] resulted from more negative amplitudes of FRN for misses [4.133±0.372, M±SE] as compared to gains [6.337±0.611, M±SE]. A main effect of *Probability* [*F*_(1,40)_ = 42.099, *p* < 0.001, η^2^_*p*_ = 0.513] reflected larger FRN for expected outcomes (4.423±0.471) as compared to unexpected outcomes (6.047±0.477) (Fig. 7). Importantly, there was a significant two-way interaction of *Valence* × *Probability* [*F*_(1, 40)_ = 10.540, *p* = 0.002, η^2^_*p*_ = 0.209]: the effect of probability for misses was smaller [*F*_(1, 40)_ = 5.294, *p* = 0.027, η^2^_*p*_ = 0.117], than for gains [*F*_(1, 40)_ = 38.101, *p* < 0.001, η^2^_*p*_ = 0.494].

We also observed a significant interaction of *Electrode* × *Probability* [*F*_(2, 68)_ = 5.275, *p* = 0.011, η^2^_*p*_ = 0.117] indicating the frontocentral maximum of the effect. The main effect of Group was not significant and no significant interactions with other factors were observed.

We further tested the effect of magnitude on FRN amplitudes. We observed a significant main effect of *Electrode* [*F*_(1,56)_ = 6.089, *p* = 0.009, η^2^_*p*_ = 0.132] supporting a frontocentral maximum of FRN. Main effect of *Valence* [*F*_(1,40)_ = 11.248, *p* = 0.002, η^2^_*p*_ = 0.219] resulted from the more negative amplitude of FRN to misses (4.292±0.381) as compared to gains (5.772±0.579). We did not observe any significant main effects of *Magnitude* and *Group*, or interaction thereof.

Analysis of dFRN (Fig. 5) showed the significant main effect of dFRN type [*F*_(2, 76)_ = 12.515, *p* < 0.001, η^2^_*p*_ = 0.238], indicating the larger *probability dFRN* (-2.454±0.528), than *valence dFRN* (-1.847±0.440) and nearly absent *magnitude dFRN* (0.691±0.518). Main effect of *Group* on dFRN was not significant, as well as *dFRN type* × *Group* interaction. Topographies clearly showed frontocentral dFRN distribution for valence dFRN and probability dFRN but not for magnitude dFRN.

**Figure 5.**
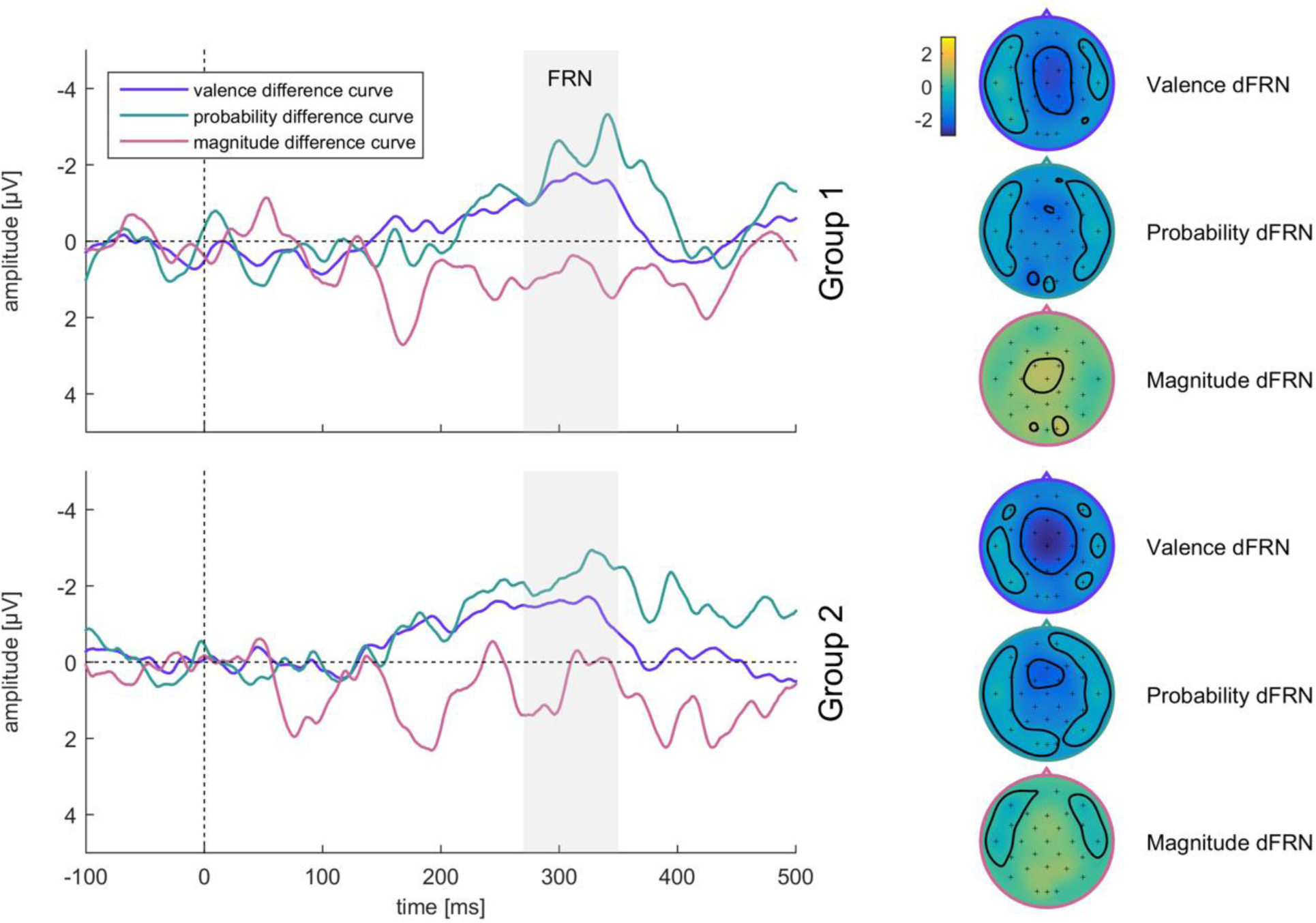
Grand-averaged visual ERP difference waveforms superimposed for three types of FRN (dFRN), calculated separately for valence (misses - gains), for probability ((unexpected misses – unexpected gains) – (expected misses – expected gains)) and for magnitude ((big misses – big gains) – (small misses – small gains)). Difference waveforms were calculated separately for two experimental groups: Group 1 – top left picture, Group 2 – bottom left picture. Time windows (270 – 350 ms) indicated by gray shading were used for individual peak amplitude measurement and for topographies calculation. Scalp topographies (right row) show dFRN distribution for valence, probability and magnitude.

### 3.3. Electrophysiological results: comparison of oddball session 1 and 2

We compared grand averaged difference waveforms obtained in the oddball sessions 1 (before MID task, baseline session) and oddball session 2 (after MID task). Fig. 6 shows superimposed grandaverage difference waveforms for auditory ERP recorded from Fz during the oddball sessions 1 and 2. ERPs were obtained for four different types of deviants separately for group 1 and group 2. In difference waveforms for all groups two main components can be seen: a negatively displaced MMN peaking at about 120 ms after stimulus onset, and a consecutive positively displaced deflection at about 230 ms, the P3a (Fig. 6). Only P3a component, and not MMN, showed a bigger amplitude during oddball session 2 as compared to session 1 in all experimental conditions.

**Figure 6.**
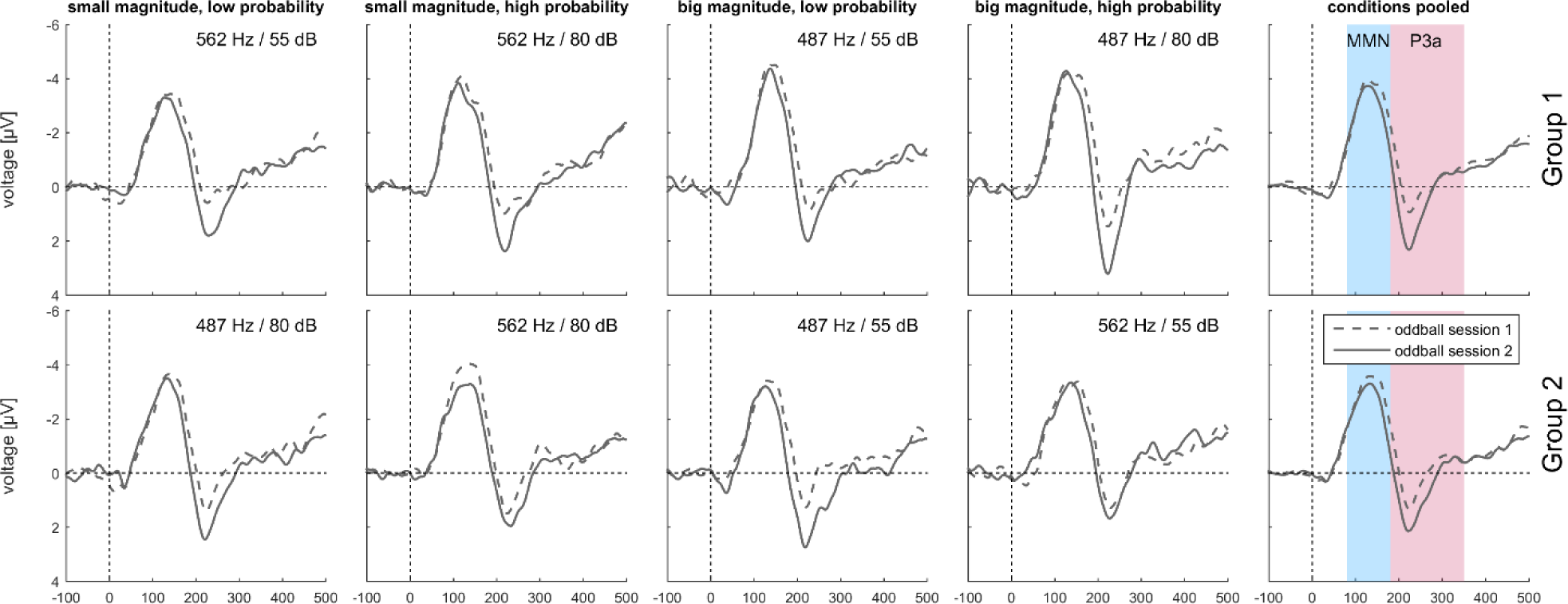
Grand-averaged deviant-minus-standard difference waveforms superimposed for two oddball sessions: before- and after-MID. Data are presented for two experimental groups: Group 1 – top row, Group 2 – bottom row. ERPs are presented for all 4 combinations of magnitude and probability of gain, constituting EV and reflected in physical parameters of the stimuli. The fifth column represents difference waveforms, derived by pooling all four conditions.

A five-way analysis of variance yielded a main effect of Session for MMN amplitudes [*F*_(1, 40)_ = 13.442, *p* = 0.001, η^2^_*p*_ = 0.252], indicating overall decrease in MMN amplitudes on the second experimental day (-1.960±0.221) compared to first experimental day (-2.398±0.220). The main effect of Electrode was also significant [*F*_(1, 53)_ = 67.662, *p* < 0.001, η^2^_*p*_ = 0.628], reflecting bigger MMN amplitudes over frontal areas. Main effect of the Group, Magnitude and Probability was not significant.

There was a significant four-way interaction Session × Magnitude × Probability × Electrode [*F*_(2, 75)_ = 4.696, *p* = 0.013, η^2^_*p*_ = 0.105], indicating decrease of MMN amplitude in condition with small magnitude and high probability on the second day (-0.724±0.212) as compared to first (-1.581±0.216).

As for the P3a amplitudes, there was a significant main effect for Session, [*F*_(1,40)_= 24.732, *p* <0.001, η^2^_*p*_ = 0.382], indicating increase in P3a amplitudes on the second experimental day, after performing MID-task (0.838±0.152) compared to baseline oddball session (0.165±0.137). The main effect of Electrode was also significant, [*F*_(2,80)_= 14.704, *p* < 0.001, η^2^_*p*_ = 0.269], due to larger P3a amplitudes in frontal areas. Main effects of Magnitude, Probability and Group were non-significant.

Given these findings, for subsequent investigation of statistical connection between electrophysiological underpinnings of reinforcement learning and plasticity of sensory input processing, we merged difference waveforms obtained for four types of deviants. Resulting waveforms are presented on Fig. 6. The frontocentral distribution of the negative and positive waves and its latency clearly suggest the activation of MMN and P3a generators (Fig. 7). MMN is a negative component, thus yellow area on difference topographies indicates overall decrease in MMN on second day. P3a is a positive component, thus yellow frontocentral spot on difference topographies shows overall increase P3a on the second day.

**Figure 7.**
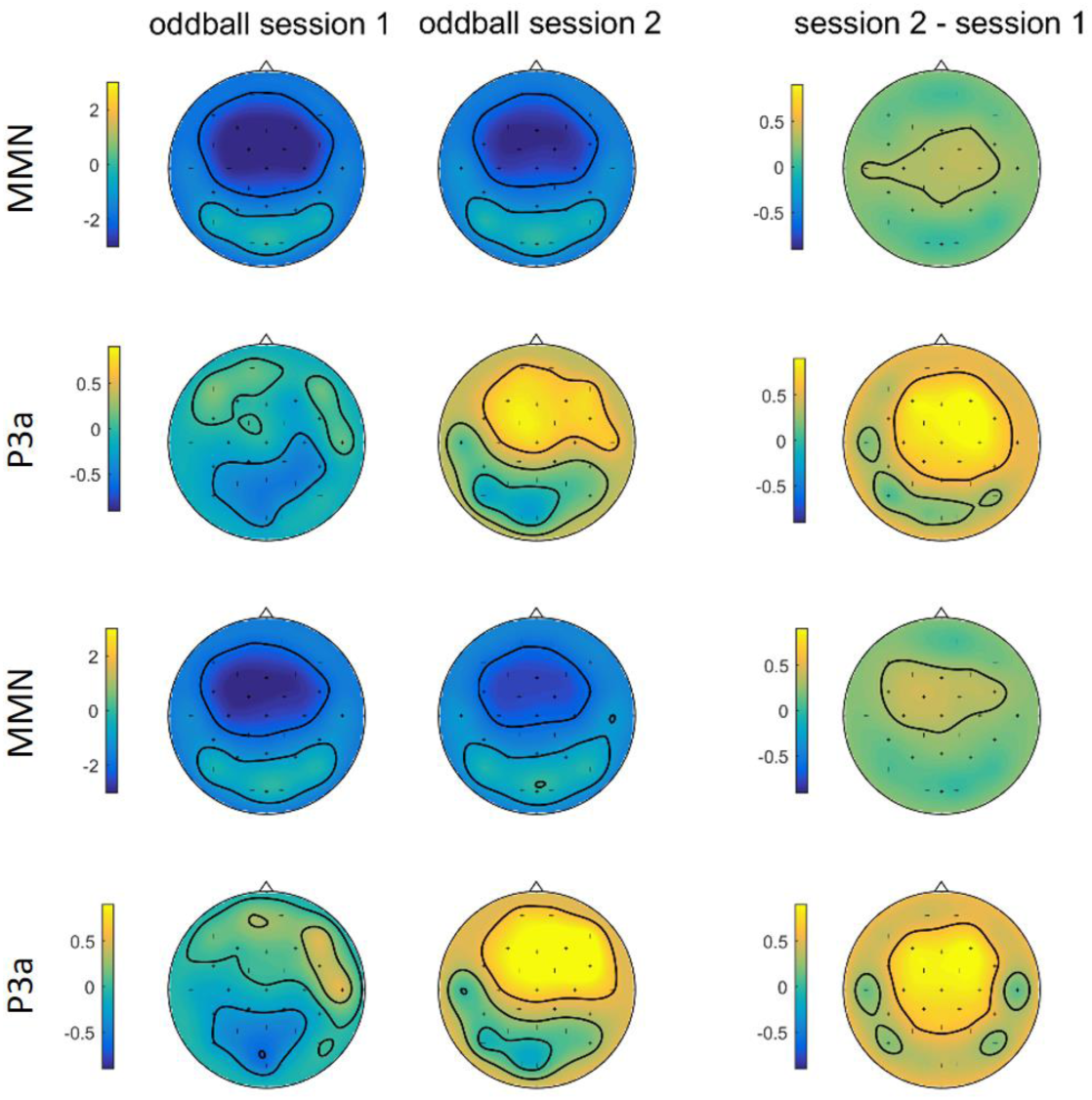
Scalp topography of the MMN and P3a during oddball session one, two, and difference topographies. Topographic maps of mean amplitude between 80 and 180 ms (MMN, first and third raw), and 180 and 350 ms (P3a, second and fourth raw) for the first oddball session (left) and second oddball session (middle), obtained for pooled condition. Difference between topoplots across two oddball session is presented on the right part of plot.

### 3.4. Results of correlational analysis

We computed correlation between changes in MMN (Cz) and P3a (Fz) amplitude across oddball sessions and dFRN, obtained for three types of difference waveforms (Fz during MID tasks). The results are shown in Fig 8.

**Figure 8.**
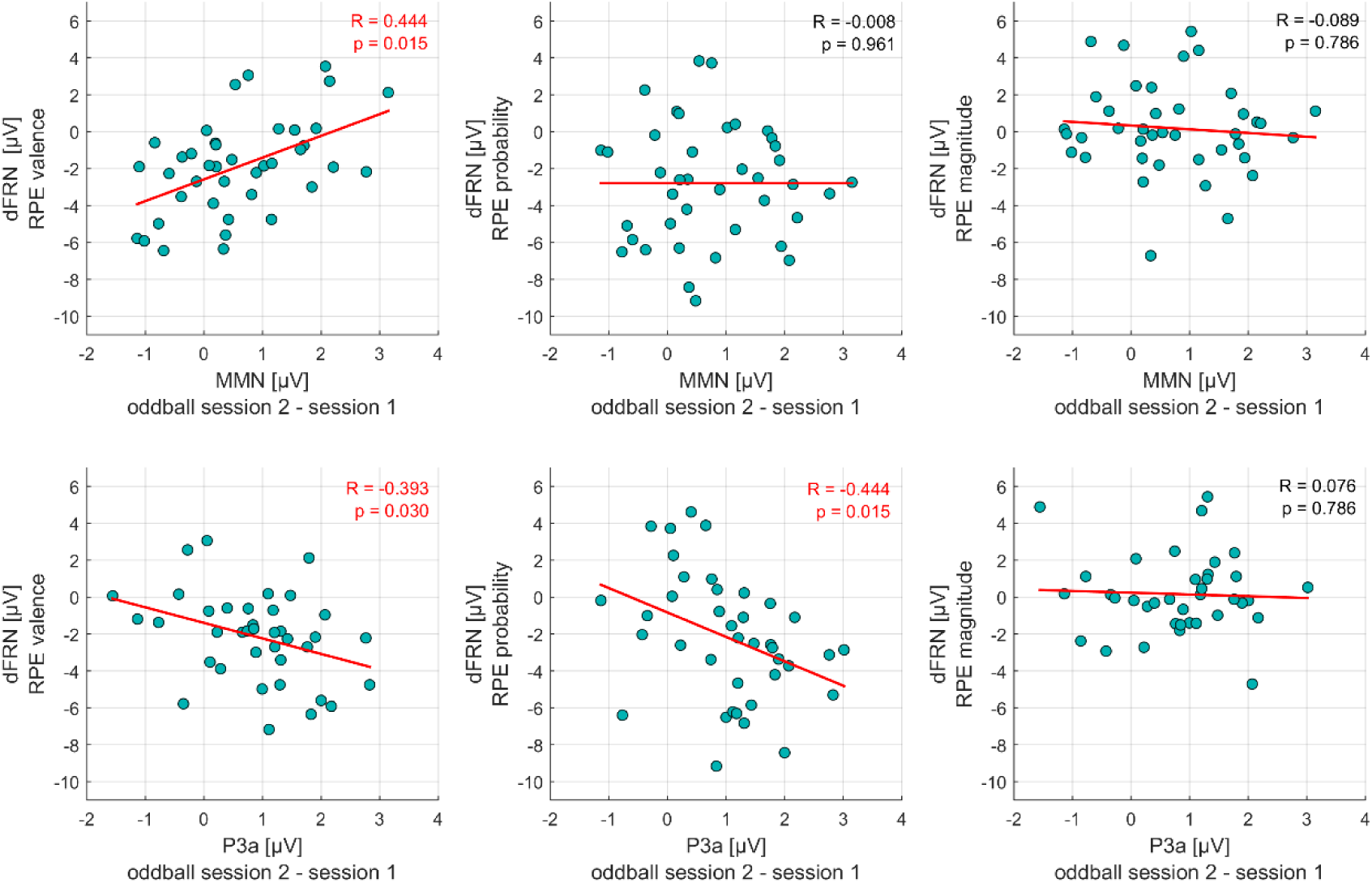
Changes in MMN (Cz, top row) and P3a (Fz, bottom row) components (oddball session 2 minus oddball session 1) as a function of valence dFRN (left), probability dFRN (middle) and magnitude dFRN (right).

Correlational analyses showed a significant positive correlation between difference MMN and valence dFRN (R = 0.444, p=0.005). Difference P3a was negatively correlated to valence dFRN (R = −0.393, p=0.015) and to probability dFRN (R = −0.444, p=0.005). However, correlation between difference MMN and probability and magnitude dFRNs, as well as difference P3a and magnitude dFRN did not reach significance level (both ps > 0.10).

Note that the significant correlations between the difference MMN and valence dFRN was negative, indicating that *MMN enlargement* in the second oddball session was significantly associated with a *bigger* valence dFRN during MID tasks. Meanwhile, correlations between P3a and valence and probability dFRNs were negative, indicating that P3a enlargement was significantly associated with more pronounced valence and probability dFRNs during MID tasks.

## 4. Discussion

Our investigation assessed the association of changes in auditory cues perception, reflected in MMN and P3a amplitudes, with individual effectiveness of learning, reflected in dFRN, elicited by feedback presentation in MID task.

Firstly, we conducted analysis of FRN and dFRN amplitudes to evaluate sensitivity of these components to valence of outcome and EV components. Our results are strongly in accordance with previous investigations, where FRN sensitivity to outcome valence was well established (Hajcak et al., 2006; Nieuwenhuis et al., 2004; Yeung and Sanfey, 2004). In our study, FRN amplitude was overall higher in misses compared to gains, for both waveforms pooled by probability and magnitude. Previously it has been proposed, that FRN reflects evaluation of positive vs. negative outcome and this binary evaluation is more pronounced in processing utilitarian (gain vs. miss) than in performance (correct vs. incorrect) feedback information (Nieuwenhuis et al., 2005). In our study the feedback in MID task comprised of both utilitarian and performance information, which ensured significant difference in processing of gains and misses. In addition to examining the effect of reward valence on the FRN amplitude, we analyzed dependence of FRN on probability and magnitude of obtained outcome. Separate pooling waveforms for feedbacks with different likelihood and magnitude allowed us to disentangle differential effect of these two RPE modulators on outcomes of different valence.

The result pattern observed for FRN amplitude modulation by probability was consistent with previous findings: probability of obtained outcome had the stronger influence on FRN amplitude, as compared to magnitude. Previous investigations demonstrated gain/loss asymmetry of probability modulation: likelihood of outcome affects waveforms for gains more strongly than for losses or omission of gain (for review, see Walsh and Anderson, 2012). This preferential sensitivity of FRN to changes in size of positive RPE might indicate different neural mechanisms underlying feedback processing in wins and in losses (Cohen et al., 2007). The design of the current study could also impact to strong influence of probability on FRN magnitude. Results of RT analysis suggest that information on probability of obtaining gain was more salient for participants than gain magnitude, as it was carrying information how fast the participant should be. This behavioral saliency together with the fact, that participant could distinguish short and long target exposition time during the course of trial, could lead to better formation of expectations.

In contrast to strong probability effect, we did not find any significant modulation of FRN amplitudes by magnitude, and magnitude-dFRN did not have pronounced negative deflection in the time window of interest. This is not a surprising finding, since the modulation of FRN by expected outcome magnitude is still under debate. In some investigations it has been postulated that FRN is not influenced by reward magnitude (Cui et al., 2013; Hajcak et al., 2003, 2006; Holroyd et al., 2006; Marco-Pallares et al., 2008; Nieuwenhuis et al., 2004; De Pascalis et al., 2010; Yeung and Sanfey, 2004), however, there is also a mounting evidence that FRN encode magnitude in addition to probability and valence (Bellebaum et al., 2010; Kreussel et al., 2012; Toyomaki and Murohashi,2005). Therefore, our results replicated previous findings of absence of FRN modulation by reward magnitude, which might be explained by relatively low difference in magnitude of potential outcome. Possibly, the difference between 4 and 20 rub was not salient enough to result in significantly distinct FRN amplitudes. In study by Bellebaum et al. (2010) it has been proposed that FRN modulation by magnitude can be clearly seen comparing the difference between big gains vs. omission of big gains and small gains vs. omission of small gains, rather then big gains with small gains. This analysis design is very similar to one that we used for magnitude-pooled ERPs. Most likely, if we had introduced bigger difference in outcome magnitudes, we would have observed results similar to Bellebaum et al. (2010).

Overall, the present study indicates that only variations in the probability of positive prediction errors affected FRN amplitudes. These results support previous assumptions, that separate neural systems perform estimation of reward probability and magnitude, and that the FRN is more sensitive to the former but not the latter component of expected value.

Our electrophysiological results are supported by modulation of RT in different types of MID probes. We found that both magnitude and probability of possible outcome strongly influenced reaction times of participants, independently from the physical parameters characterizing the sounds implemented as incentive cue. This indicates that participants of both groups correctly integrated the information encoded in the incentive cue to consequently adjust their behavior, i.e. expressing higher latency in response to high probability and low magnitude trials. Absence of significant interaction between magnitude and probability might be explained by the separate influence of two components of EV on RTs, but not EV itself. Previous investigations, where magnitude and/or probability of outcome in MID task signaled by incentive cue have been manipulated, showed that factors *Magnitude* and *Probability* similarly influenced reaction times of participants (Helfinstein et al., 2013; Knutson et al., 2003, 2005; Rademacher et al., 2014). In agreement with previous finding with analogous implementation of probability encoding (Knutson et al., 2005) we found stronger probability modulatory effects on RTs as compared with magnitude. Due to association of probability of prospective gain to the duration of target exposition in MID, we can assume that identification of probability might be more critical than magnitude for good performance.

To investigate the experience-induced plasticity as a function of the magnitude and probability, we focused on changes in MMN and P3a components. Across the two consecutive oddball sessions we could identify a decrease in the amplitude of the MMN component and an increase in the amplitude of the P3a component.

Decrease in MMN amplitude, registered for some participants, might be interpreted as lack of evoked plastic changes in auditory cortex, indicating absence of improvement in preattentive stimuli discrimination (Gurevicius et al., 2013), whereas increase is believed to associated with evoked plastic changes (Atienza et al., 2002; Näätänen, 2008a). Some studies find no association of MMN changes with performance (Näätänen et al., 1993a; Uther et al., 2006). As was proposed by (Näätänen et al., 1993b), in human participants with good initial discrimination of auditory stimuli, the MMN amplitude is already large and cannot increase during the course of discrimination training. Taken together, this and other MMN findings associated with plastic changes (Novak et al., 1990; Tiitinen et al., 1997), suppose that decrease in MMN amplitude across two oddball sessions could be related to initially good discrimination of frequency and intensity of deviant sounds across participants (Novak et al., 1990; Tiitinen et al., 1997). We consistently observe a decrease in MMN amplitude across two oddball sessions, as expected by the high discrimination performance of frequency and intensity of deviant sounds across participants. Notably, the overall MMN decrease from the first to the second oddball was replicated in both experimental groups.

We observed a significant increase in P3a amplitude in the second oddball session across all four EVs. We hypothesized that the experience-induced P3a enhancement is associated with the assigning EVs to auditory tones during MID task and represent the involuntary attention switch towards those tones (Escera and Corral, 2007; Escera et al., 1998). This experience-induced plasticity might be mediated by top-down process, resulting in the enhanced change detection (Seppänen et al., 2012), which, in its turn, facilitates processing of auditory stimuli.

The absence of prominent patterns in the amplitude change of MMN or P3a for different levels of probability and magnitude, or their specific combination, constituting EV, can be explained by equal saliency of tones for MID task performance. Alternatively, these changes could be registered only because of adaptation effect, because of repeated exposure to tones in oddball and MID task. Due to the similarity of changes in MMN and P3a in different auditory tones, we pooled together the four incentive cues for further investigation of individual differences in plastic changes.

In previous studies it has been proposed that size of FRN predicts effectiveness of learning (Luft 2014). The majority of these studies, which found evidence in support of this hypothesis, used paradigms requiring building probabilistic associations rather than error-based learning (Arbel et al., 2013; Cohen and Ranganath, 2007; Frank et al., 2005; Van Der Helden et al., 2010; Philiastides et al., 2010; Yasuda et al., 2004; for review, see Luft, 2014). Here we tested whether components associated with processing of RPE sign (valence dFRN) and size (probability and magnitude dFRN) are predictive of plastic changes in sensory stimuli perception.

MMN amplitude enlargement after MID task correlated with valence dFRN component amplitude but not with probability or magnitude dFRNs. This suggests that participants demonstrating enlargement of MMN after two days of training, also had pronounced valence – dFRN, reflecting RPE processing, while participants, that had small or absent valence dFRN showed decrease in MMN amplitude on the second day. The larger MMN amplitudes, registered on the second day in subjects with bigger valence dFRN, might indicate an induced plasticity of auditory cortex, driven by MID task performance (Gottselig et al., 2004; Näätänen, 2008b; Pantev and Herholz, 2011).

Notably, we found an association between experience-induced P3a amplitude changes and probability dFRN, and not valence or magnitude dFRN. This showed that participants with pronounced probability – dFRN have enlargement of P3a recorded in oddball after the MID task performance. Probability dFRN reflects not only difference in processing of gains and misses, but also carries information about processing of RPEs of different size. Difference in the processing of small and big RPEs would emerge only if participant had expectations about probability of a particular type of feedback, which in the case of our paradigm implies that the participant took into account information preceding feedback presentation. Information about probability of prospective gain can be derived both from physical characteristics of the auditory cue and from duration of target presentation itself. But in both cases indication of probability was not so explicit as in case of visual cues in classical MID task (Knutson et al., 2005), so we suppose, that only highly motivated and emotionally engaged participants payed attention to these parameters and successfully formed expectations. Thus, involuntary attention switch, as reflected by P3a changes, was associated with differential processing of RPEs of different size.

Moreover valence dFRN and probability dFRN were associated with changes in different steps of perceptual processing of auditory cues. These findings might be an indirect evidence of involvement of different neural pathways in processing of feedback valence and likelihood (degree of violation of expectations) (Ferdinand and Opitz, 2014a). It has been previously shown that while valence and magnitude modulated BOLD signal in nucleus accumbens, probability affected signal coming from medial prefrontal cortex (mPFC), and ACC was proposed as a hub, integrating information about valence, probability and magnitude (Knutson et al., 2005). In other studies the difference in processing gains and losses or omission of gain was also localized in ventral striatum (Ferdinand and Opitz, 2014b; Satterthwaite et al., 2012; Späti et al., 2015; Zink et al., 2004). Involvement of mPFC in probability estimation is well described in literature (Alexander and Brown, 2012; Brown and Braver, 2005, 2007; Krawitz et al., 2011).

In recent meta - analysis it has been shown, that effects of probability and magnitude on dFRN amplitude have a later time course compared to valence. This can be explained by influence of P300 component, following the FRN and known to be associated with saliency processing (Sambrook and Goslin, 2015a). Interestingly, in the same meta-analysis, mentioned above, the probability and magnitude dFRN was stronger, when participants had control over outcome, i.e. to obtain gain, they implemented known rule (Sambrook and Goslin, 2015a). Authors assume, that this feeling of agency enhanced saliency of the feedback and this might reflect that dFRN is rather a part of instrumental conditioning system.

Taken together, the evidence of different neural underpinnings of valence and likelihood processing (Shizgal, 1997) and the assumption that multiple components operate within the FRN temporal interval could clarify the differential pattern of correlation that we observed. We can only speculate about dependency of plastic changes on the size of FRN, because our design does not address causality. However, we can assume that basic evaluation of positive vs. negative outcome related to changes in early stages of perception of relevant auditory stimuli. Whereas processing of feedback saliency is associated with future enhancement of attention redirection towards these stimuli.

## 5. Conclusion

Considered in the framework of the RL theory of the FRN (Holroyd and Coles, 2002) our findings support the hypothesis that the degree of plastic changes in auditory cues perception depend on the strengths of reinforcement signal. Overall, our results showed that continuing valuation of auditory cues associated with different EVs evokes plastic changes associated with enhanced involuntary attention switch. Observed signatures of neuro-plasticity of the auditory cortex may play an important role in learning and decision-making processes through facilitation of perceptual discrimination of valuable external stimuli. Thus, the sensory processing of options and the evaluation of options are not independent as implicitly assumed by the dominant neuroeconomics models of decision-making.

## 6. Acknowledgements

The study was supported by the grant 16-18-00065 of the Russian Science Foundation.

